# Inactivation of SARS Coronavirus 2 and COVID-19 patient samples for contemporary immunology and metabolomics studies

**DOI:** 10.1101/2021.10.22.465481

**Authors:** Devon J. Eddins, Leda Bassit, Joshua D. Chandler, Natalie S. Haddad, Katie L. Musall, Junkai Yang, Astrid Kosters, Brian S. Dobosh, Mindy R. Hernández, Richard P. Ramonell, Rabindra M. Tirouvanziam, F. Eun-Hyung Lee, Keivan Zandi, Raymond F. Schinazi, Eliver E.B. Ghosn

## Abstract

In late 2019, severe acute respiratory syndrome coronavirus 2 (SARS-CoV-2) emerged from Wuhan, China spurring the Coronavirus Disease-19 (COVID-19) pandemic that has resulted in over 219 million confirmed cases and nearly 4.6 million deaths worldwide. Intensive research efforts ensued to constrain SARS-CoV-2 and reduce COVID-19 disease burden. Due to the severity of this disease, the US Centers for Disease Control and Prevention (CDC) and World Health Organization (WHO) recommend that manipulation of active viral cultures of SARS-CoV-2 and respiratory secretions from COVID-19 patients be performed in biosafety level 3 (BSL3) containment laboratories. Therefore, it is imperative to develop viral inactivation procedures that permit samples to be transferred and manipulated at lower containment levels (i.e., BSL2), and maintain the fidelity of downstream assays to expedite the development of medical countermeasures (MCMs). We demonstrate optimal conditions for complete viral inactivation following fixation of infected cells with paraformaldehyde solution or other commonly-used branded reagents for flow cytometry, UVC inactivation in sera and respiratory secretions for protein and antibody detection assays, heat inactivation following cDNA amplification of single-cell emulsions for droplet-based single-cell mRNA sequencing applications, and extraction with an organic solvent for metabolomic studies. Thus, we provide a suite of protocols for viral inactivation of SARS-CoV-2 and COVID-19 patient samples for downstream contemporary immunology assays that facilitate sample transfer to BSL2, providing a conceptual framework for rapid initiation of high-fidelity research as the COVID-19 pandemic continues.

## 1. Introduction

At the end of 2019, a novel betacoronavirus, SARS-CoV-2 (1), emerged from Wuhan in the Hubei province of China causing viral pneumonia that progressed to severe or critical disease in ~20% of infected patients, where in the most critical cases, patients would present with respiratory failure and require mechanical ventilator support in the intensive care unit (2–4). Since then, much research effort has been focused on better understanding pathogenesis and immunity to SARS-CoV-2 (5). However, due to the severity of disease, work with infectious patient samples (primarily samples from the airways) and active viral cultures require biosafety level 3 (BSL3) containment facilities as with other closely related betacoronaviruses including SARS-CoV and Middle Eastern Respiratory Syndrome–Coronavirus (MERS-CoV) (1, 6, 7). This can restrict research activity where containment facilities are not available. As the COVID-19 pandemic continues to surge with emergence of new variants (8), there is a continued need to conduct frontier research on COVID-19 immunology and pathogenesis to develop and refine medical counter measures (MCMs) to protect at risk populations and those disproportionately affected by COVID-19 disease (9–11). To facilitate this work, it is imperative to understand effective viral inactivation protocols that have minimal effects on assay readouts.

Though SARS-CoV-2 shares ~80% sequence homology with SARS-CoV (1), it is essential to evaluate efficacy of existing inactivation procedures on novel, independent viral strains. For example, MERS-CoV is also closely related to SARS-CoV but there are notable differences in the inactivation efficacy using gamma-irradiation between the viruses (12, 13). Indeed, there have been earlier reports of efficient viral propagation and inactivation procedures for SARS-CoV-2 using heat, fixatives/chemicals/surfactants (e.g., formaldehyde/Trizol^®^/Triton X-100), and UVC irradiation (14–19), which are comparable to SARS-CoV and MERS-CoV (12, 13, 20–22). However, though most of these studies highlight efficient and effective viral inactivation protocols (15–19), the effect of inactivation on the fidelity of downstream assay readouts and analysis remain largely unknown. Specifically, a detailed report on viral inactivation protocols and how they influence contemporary immunology assays is notably lacking. Contemporary immunological assays including ELISA/Luminex/Mesoscale assays for antibody/protein detection (9, 23–25), metabolomics (26, 27), high-dimensional (Hi-D) flow cytometry (9, 28–30), and multi-omics single-cell mRNA sequencing (scRNA-seq) (9, 31, 32) are vital resources to develop and evaluate MCMs for the ongoing COVID-19 pandemic.

To address the need to successfully inactivate virus and permit transfer of material from BSL3 to a lower containment (i.e., BSL2) environment for high-fidelity downstream assays, we examined the efficiency of several viral inactivation methods for contemporary immunological assays including flow cytometry, serology/protein detection, scRNA-seq, and high-throughput metabolomic experiments using both culture-derived virus and infected respiratory samples from COVID-19 patients. Here, we report complete viral inactivation following fixation with 4% PFA or 1.6X BD FACS™ Lysis Solution (~2.5% formaldehyde and ~8.3% diethylene glycol) for 30 min at room temperature for flow cytometry, UVC inactivation at ~4,000 μwatts/cm^2^ for 30 min in sera and respiratory secretions for protein/antibody detection assays, heat inactivation following first-round cDNA amplification of single-cell emulsions for droplet-based scRNA-seq, and metabolite extraction for 30 min with 4 volumes of a 1:1 acetonitrile:methanol solution with 12.5 μM/L D5-benzoylhippuric acid at 4°C for metabolomic studies. These results will serve as conceptual framework and promote rapid initiation of cutting-edge immunology studies as the COVID-19 pandemic continues to evolve and for other risk group 3 agents that require higher containment.

## 2. Methods

### 2.1 Ethics & Biosafety statements

COVID-19+ patients were recruited from the Intensive Care Units of Emory University, Emory St. Joseph’s, Emory Decatur, and Emory Midtown Hospitals (severe) or the Emory Acute Respiratory Clinic (mild), and healthy adults were recruited from the Emory University Hospital. All studies were approved by the Emory Institutional Review Board (IRB) under protocol numbers IRB00058507, IRB00057983 and IRB00058271. Informed consent was obtained from the patients when they had decision making ability or from a legal authorized representative (LAR) if the patient was unable to provide consent. We collected both blood and non-induced sputum (healthy/mild) or endotracheal aspirate (ETA; severe). Study inclusion criteria included a confirmed COVID-19 diagnosis by PCR amplification of SARS-CoV-2 viral (v)RNA obtained from naso-/oro-pharyngeal swabs, age of 18 years or greater, and willingness and ability to provide informed consent. All work with infectious virus and respiratory samples from COVID-19 patients was conducted inside a biosafety cabinet within the Emory Health and Safety Office (EHSO)- and United States Department of Agriculture (USDA)-approved BSL3 containment facility in the Health Sciences Research Building at Emory University following protocols approved by the Institutional Biosafety Committee (IBC) and Biosafety Officer.

### 2.2 Virus and cells

African green monkey (*Cercopithecus aethiops*) kidney epithelial cells (Vero E6 cells; ATCC^®^ CRL-1586™) were maintained in complete (c)DMEM containing: 1X DMEM supplemented with 25 mM HEPES, 2 mM L-glutamine,1 mM sodium pyruvate, 1X non-essential amino acids (NEAA), 1X antibiotic/antimycotic solution (all from Corning) and 10% heat-inactivated FBS (Gibco), unless indicated otherwise. Human lung adenocarcinoma epithelial cells (Calu-3 cells; ATCC^®^ HTB-55™) were maintained in cMEM containing: 1X MEM (Corning) supplemented with 1X antibiotic/antimycotic solution and 10% heat-inactivated FBS unless indicated otherwise. Primary leukocytes from the airways of severe COVID-19 patients were collected bedside via endotracheal aspiration (ETA) and whole blood collected by standard venipuncture, then processed as previously described (9). SARS-CoV-2 USA-WA1/2020 (hereafter SCV2-WA1) was provided by BEI Resources (Manassas, VA, USA). Virus was propagated in Vero E6 cells as previously described (16, 33) and titer determined by TCID_50_ (TCID_50_/mL) or plaque assays (PFU/mL). Low-passage (P1 or P2) virus stocks were used throughout this study.

### 2.3 Infectivity assays

#### 2.3.1 Plaque Assays with Methylcellulose

Vero E6 cells were seeded in 6-well plates (Falcon) with 5 × 10^5^ cells/well in 5% DMEM 24 h prior to infection and checked to verify ≥80% confluency. 10-fold dilutions of virus, respiratory secretions, and/or scRNA-seq emulsion in serum-free DMEM (200 μL) were incubated on Vero E6 monolayers for 1 h absorption at 37°C with rocking at 15 min intervals. After absorption, cells were overlain with 2% methylcellulose (MilliporeSigma) in 2% DMEM for 72 h at 37°C in a 5% CO_2_, humidified incubator. 72 h post-infection (hpi), methylcellulose was carefully removed, and cells gently rinsed once with 1X HBSS (Corning). Monolayers were fixed and plaques visualized with a solution of 0.4% crystal violet by weight in 80% methanol (MilliporeSigma) and 4% PFA (Electron Microscopy Sciences) for 20 min at room-temperature (RT).

#### 2.3.2 Plaque Assays with Agarose

Vero E6 cells were seeded in 6-well plates with 5 × 10^5^ cells/well in 5% DMEM 24 h prior to infection and checked to verify ≥80% confluency. 10-fold dilutions of virus, respiratory secretions, and/or scRNA-seq emulsion in serum-free DMEM (200 μL) were incubated on Vero E6 monolayers for 1 h absorption at 37°C with rocking at 15 min intervals. After absorption, cells were overlain with 2 mL 0.5% immunodiffuse agarose (MP Biomedicals) in 1X DMEM supplemented with 5% FBS, 2 mM L-glutamine,1 mM sodium pyruvate, 1X NEAA, 1X sodium bicarbonate, and 1X antibiotic/antimycotic solution. 72 hpi, a second 2 mL overlay of 0.5% immunodiffuse agarose in a 1X HBSS solution with 0.026% neutral red (MilliporeSigma) was added for ≥3 h to visualize plaques.

#### 2.3.3 TCID_50_ assays

Vero E6 cells were seeded in 96-well plates with 2 × 10^4^ cells/well in 5% MEM 24 h prior to infection and checked to verify ≥80% confluency. 10-fold dilutions of stock SCV2-WA1 virus in serum-free MEM (100 μL) were incubated on Vero E6 monolayers in quadruplicates for 2 h absorption at 37°C without rocking. Following absorption, the inoculum was removed, and cells cultured in 2% MEM. Cells were assessed daily for cytopathic effect (CPE) compared to mock-infected negative controls by microscopy for 6 d. Calculations for 50% tissue culture infectious dose (TCID_50_) were performed using either the Spearman-Käber (34) or Reed and Muench (35) methods as previously described (36).

#### 2.3.4 Focus Reduction Neutralization Assays (FRNA)

Vero E6 cells were seeded in 96-well plates with 2 × 10^4^ cells/well in 5% DMEM 24 h prior to infection and checked to verify ≥80% confluency. Dilutions of virus and/or virus treated with inactivation reagents in Opti-MEM™ (50 μL) were incubated on Vero E6 monolayers for 2 h absorption at 37°C without rocking. After absorption, cells were overlain with 2% methylcellulose in Opti-MEM™ (Gibco) supplemented with 2% FBS, 2.5 μg amphotericin B (MilliporeSigma), and 20 μg/mL ciprofloxacin (MilliporeSigma) for 72 h at 37°C in a 5% CO_2_, humidified incubator. 72 hpi, methylcellulose was carefully removed, and the cells fixed with 1:1 methanol/acetone mixture for 30 min at RT, then blocked with 200 μL 5% milk in 1X PBS for 20 min. Cells were incubated with an anti-SARS-CoV-2 spike RBD polyclonal antibody (Gentaur) at 1:3000 overnight at 37°C. Cells were washed to remove excess antibody, then incubated with a secondary HRP-conjugated anti-human IgG for 1 h at 37°C. Cells were washed to remove excess antibody and foci visualized using the TrueBlue™ Peroxidase Substrate (SeraCare Life Sciences) incubated for 1 h at RT with rocking prior to imaging with an ELISpot reader for foci quantification. Data reported as focus forming units per mL (FFU/mL).

### 2.4 Inactivation by fixative solutions

Vero E6 cells were infected at a multiplicity of infection (MOI) of 0.01 and cultured in 6-well plates for 48 h with cMEM or cells from ETAs of COVID-19-infected patients were fixed with a freshly prepared 2% or 4% PFA solution (20% stock diluted in 1X HBSS; Electron Microscopy Sciences) or a 1.6X BD FACS™ Lysis Solution (1:6 in sterile dH_2_O; BD Biosciences) for the indicated time points at RT. Unfixed cells were incubated with 1X HBSS. Following fixation, cells from each condition were washed twice and resuspended in cMEM for an additional 48 h incubation at 37°C in a humidified, 5% CO_2_ incubator. Culture supernatants were then collected and either plated immediately or frozen at −80°C for analysis by plaque assay.

### 2.5 Inactivation by ultraviolet C (UVC) radiation

50-1000 μL aliquots of SCV2-WA1 virus stock (2.1 × 10^5^ PFU/mL), respiratory supernatant (additional samples combined with our previously published data (9)), or patient sera were collected for UV inactivation. Samples were positioned 2-3 cm from the light source and exposed to 254 nm UVC light at maximum intensity (~4000 μwatts/cm^2^) for 30 min using a Spectrolinker™ XL-1000 UV crosslinker (Spectronics Corporation) in either clear 2 mL microcentrifuge tubes (positioned on their side) or a 96-well plate (with the lid removed). Samples were either plated immediately or frozen at −80°C for analysis by plaque assay.

### 2.6 Inactivation for metabolomic assays

To assess a nontoxic concentration of the metabolite extraction solvent (50% acetonitrile, 50% methanol, and 12.5 μM/L D5-benzoylhippuric acid) for subsequent FRNA, cytotoxicity tests were performed in Vero E6 cells via MTS assay using the CellTiter 96^®^ Non-Radioactive Cell Proliferation kit (Promega) as previously described (37). Uninfected Vero E6 cells were incubated with the extraction solvent, diluted in Opti-MEM™ at 1:10, 1:100, and 1:1000 in triplicate at 37°C, 5% CO_2_ for 72 h with 100, 10, and 1 μM cycloheximide as positive control. After 72 h, the MTS tetrazolium compound was added to the cells and incubated for an additional 2 h. To determine the number of viable cells in each well, the absorbance was measured at 490 nm using a 96-well plate reader (BioTek). Cytotoxicity was expressed as the dilution of the extraction solvent that inhibited cell proliferation by 50% (IC50) and calculated using the Chou and Talalay method (38). SCV2-WA1 (2.5 × 10^4^ TCID_50_/mL) was incubated with or without the extraction solvent (1:4) or Triton X-100 for 30 min at 4°C, then centrifuged at 20,000 x g for 10 min at 4°C. Supernatants were collected and diluted in Opti-MEM™ to final concentrations of 1:100 the extraction solvent or 1% Triton X-100, and >100 FFU/mL SARS-CoV-2 per well for analysis by FRNA.

### 2.7 Inactivation for scRNA-seq (10X Genomics)

SARS-CoV-2-infected Vero E6 cells (MOI 0.04 for 72 h) or Calu-3 cells (MOI 0.04 for 48 h) were encapsulated for scRNA-seq following the manufacturer’s protocol “Chromium Next GEM Single Cell V(D)J Reagent Kits v1.1 User Guide with Feature Barcode technology for Cell Surface Protein” (document number CG000208; 10X Genomics) targeting 20,000 and 10,000 cells, respectively. An aliquot of the emulsion was collected following encapsulation for analysis by plaque assay. The remaining emulsion was processed for cDNA synthesis reaction following the manufacturer’s protocol with reagent volumes adjusted to reflect the reduced reaction volume after taking aliquots for plaque assays. Polymerase chain reaction (PCR) amplification profile was 45 min at 53°C followed by 5 min at 85°C. An additional aliquot of the cDNA suspension was collected following PCR reactions and either plated immediately or frozen at −80°C for analysis by plaque assay. Plaque assays were also performed on the encapsulation emulsion alone (without including virus-infected cells) to evaluate reagent cytotoxicity on Vero E6 monolayers.

### 2.8 Luminex proteomic serology assays

Plasma from whole blood of COVID-19 patients was isolated via centrifugation at 400 x g for 10 min at 4°C. To remove platelets, the isolated plasma was centrifuged at 4,000 x g for 10 min at 4°C. Plasma samples were stored at −80°C until analyzed. Luminex serology assays were performed as previously described (24). In brief, ~50 μL of coupled microsphere mix was added to each well of 96-well clear-bottom black polystyrene microplates (Greiner Bio-One) at a concentration of 1,000 microspheres per region per well. All wash steps and dilutions were performed with 1% BSA in 1X PBS (hereafter assay buffer). Sera were assayed at 1:500 dilutions (in assay buffer) and surveyed for anti-SARS-CoV-2 N or RBD antibodies by 1 h incubation on a plate shaker at 800 rpm in the dark. Following incubation, wells were washed five times with 100 μL of assay buffer using a BioTek 405 TS plate washer, then 3 μg/mL PE-conjugated goat anti-human IgA, IgG and/or IgM (Southern Biotech) was applied. After 30 min incubation, wells were washed three times in 100 μL of assay buffer, then resuspended in 100 μL of assay buffer for acquisition and analysis using a Luminex FLEXMAP 3D instrument and xPONENT 4.3 software (Luminex). Median fluorescent intensity (MFI) using combined or individual detection antibodies (i.e., anti-IgA, anti-IgG, or anti-IgM) was measured and the background value of assay buffer was subtracted from each serum sample result to obtain MFI minus background values (net MFI).

### 2.9 Metabolomic assays

Metabolites in human plasma were extracted from the National Institute of Standards and Technology (NIST) “Standard Reference Materials 1950” after mock or UVC treatment by addition of four volumes of 50% acetonitrile, 50% methanol, and 12.5 μM/L benzoyl-D5-hippuric acid (extraction solvent). Samples were vortexed for 10 seconds, incubated on ice for 30 min, then centrifuged at 20,000 x g for 10 min at 4°C. The clear supernatant was aliquoted and injected (2.5 μL) on a Vanquish Horizon liquid chromatograph coupled to Q Exactive High Field (ThermoFisher). A 150 mm × 2.1 mm ZIC-HILIC (MilliporeSigma) column and matching guard column were used to separate polar metabolites. Metabolites were ionized in both positive and negative mode and analyzed in full scan mode (67-1000 m/z). Pooled quality control samples comprising equal proportions of every study sample were used to generate ddMS2 (Top20N) spectra of metabolites, evaluate assay reproducibility, and correct batch drift. Compound Discoverer 3.2 (ThermoFisher) was used to quantify peak areas and assign annotations based on a local library of reference standards or via matching metabolites to reference spectra in mzCloud (mzcloud.org). Data were imported to Prism 9 for graphing and statistical analyses by unpaired t-tests for untreated versus UVC-treated replicates, assuming individual variance, using the adaptive linear (two-step) step-up (Benjamini, Krieger, and Yekutieli) method (39) to control the false discovery rate (FDR), and a desired FDR (Q) of 10% for multiple comparisons.

### 2.10 scRNA-seq data alignment, dimensionality reduction, and clustering

The Cell Ranger Software (v.5.0.0; 10X Genomics) was used to perform cell barcode processing and single-cell 5′ unique molecular identifier (UMI) counting. To detect SARS-CoV-2 reads, a customized reference genome was built by integrating human GRCh38 and SARS-CoV-2 genomes (severe acute respiratory syndrome coronavirus 2 isolate Wuhan-Hu-1, complete genome, GenBank MN908947.3). Splicing-aware aligner STAR (40) was used in FASTQ alignments. Cell barcodes were then determined based on the distribution of UMI counts automatically. The filtered gene-barcode matrices were first normalized using ‘LogNormalize’ method in Seurat v.3 (41) with default parameters. The top 2,000 variable genes were then identified using the ‘vst’ method by the ‘FindVariableFeatures’ function. Principal Component Analysis (PCA) was performed using the top 2,000 variable genes, then UMAPs generated using the top 30 principal components to visualize cells. Graph-based clustering was performed on the PCA-reduced data for clustering analysis with the resolution set to 0.8 to obtain the clusters. The total viral UMIs were the sum of the UMIs of the 12 SARS-CoV-2 genes (42).

### 2.11 SARS-CoV-2 quantitative reverse transcription PCR (RT-qPCR)

Stock SCV2-WA1 virus and patient samples (400 μL) were thoroughly mixed 1:1 with 2X DNA/RNA Shield™ and incubated at RT for 20 min for inactivation and vRNA was extracted using the *Quick*-RNA™ Viral Kit (Zymo Research) following the manufacturer’s protocol. Complimentary (c)DNA was synthesized using the High-Capacity cDNA Reverse Transcription Kit (Applied Biosystems™) per manufacturer’s instructions and diluted 1:5 in nuclease-free water, then 10 μL of diluted cDNA was used with the NEB Luna Universal Probe qPCR Mastermix (New England BioLabs^®^) following the manufacturer’s protocol and RT-qPCR performed in 384-well plates using a QuantStudio™ 5 Real-Time PCR System (Applied Biosystems™). Primer/probe pairs were: AGAAGATTGGTTAGATGATGATAGT (forward primer), TTCCATCTCTAATTGAGGTTGAACC (reverse primer), and /56-FAM/TCCTCACTGCCGTCTTGTTGACCA/3IABkFQ/ (probe), which were designed from sequences previously described (43) (Integrated DNA Technologies; IDT). To generate a standard curve and quantify SARS-CoV-2 genome copies, a gBlock with the sequence: AATTAAGAACACGTCACCGCAAGAAGAAGATTGGTTAGATGATGATAGTCAACAAACTGTTGGTCAACAAGACGGCAGTGAGGACAATCAGACAACTACTATTCAAACAATTGTTGAGGTTCAACCTCAATTAGAGATGGAACTTACAGTTTCAGTGTTCAATTAA (IDT) was used as a standard.

To determine PFU equivalents (ePFU) from respiratory samples, vRNA was extracted from 10-fold serial dilutions of stock SCV2-WA1 of a known titer for RT-qPCR to generate a standard curve from which number of genome copies per PFU could be extrapolated as previously described (44). Culture supernatants from mock-(1X HBSS) and SARS-CoV-2-infected Calu-3 (MOI 0.04) cells were utilized as additional controls.

## 3. Results

### 3.1 Inactivation by fixation

To evaluate the ability of commonly-used fixatives in flow cytometry (here, formaldehyde-based and PFA) to completely inactivate SARS-CoV-2-infected cells for transfer to lower containment settings, we performed a time-course of inactivation (Fig. 1A) using 2% and 4% PFA (diluted from a 20% stock, see methods) along with 1.6X BD FACS™ Lysis Solution (10X stock diluted 1:6 in sterile dH_2_O). Cells exposed to 2% and 4% PFA for 15 min at room temperature were still able to produce infectious virus when returned to culture for 48 h (Fig. 1B,C), with a decrease in viral titer (PFU/mL). This is in contrast to a previous report that indicated 10 min treatment with 4% PFA is sufficient to inactivate virus (14). However, infectious virus was not detected by plaque assay following 15 min exposure to 1.6X BD FACS™ lysis solution and at all subsequent time points (Fig. 1B,C). Cells treated with 4% PFA for 30 min at RT were no longer infectious, however, 60 min was required to fully inactivate virus in cells treated with 2% PFA at RT. These data indicate that common fixation protocols (30 min fixation at RT) for commercially-available fixatives, specifically those with ≥4% PFA, are sufficient for inactivating SARS-CoV-2 infected cells.

**Figure 1.**
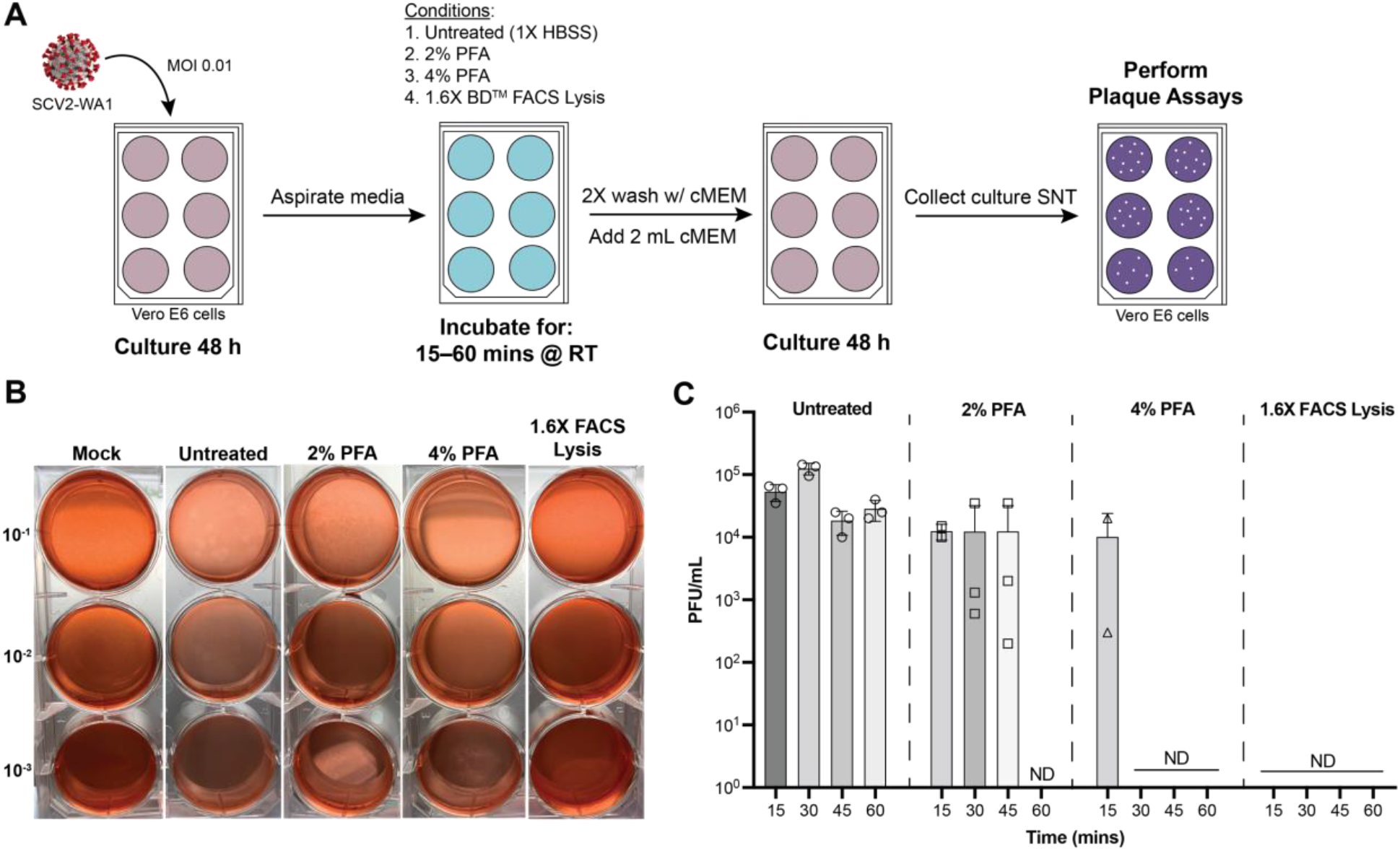
Fixation with commercially-available fixatives promote complete inactivation of SARS-CoV-2-infected cells amenable to flow cytometric analyses. A. Schematic of inactivation time course performed to evaluate inactivation efficiency. SNT = supernatant. B. Representative plaque assays from inactivation time course. C. Quantification of viral load for the 4 fixatives across the 4 time-points evaluated. ND = Not Detected (by plaque assay).

### 3.2 UVC inactivation

#### 3.2.1 Inactivation of respiratory secretions and viral stocks

Though complete inactivation of SARS-CoV viral stocks can be achieved with <15 min exposure to UVC-irradiation (12), a follow-up report for SARS-CoV inactivation in non-cellular blood products in PBS solutions recommended 40 min exposure to inactivate virus (20). Therefore, we selected 30 min exposure to UVC-irradiation at maximum intensity (~4000 μwatt/cm^2^) for both culture-derived SARS-CoV-2 viral stocks and respiratory secretions from COVID-19 patients. First, we determined viral load in the respiratory secretions by extrapolating ePFU/mL from RT-qPCR results of respiratory secretions (see methods) (44). Using a standard curve generated from virus stock of a known titer (Fig. 2A), we determined 1 PFU to be equivalent to 73 SARS-CoV-2 genome copies in RT-qPCR data. This allowed for more accurate viral detection and quantification (Fig. 2B) since endotracheal aspirate (ETA) and bronchoalveolar lavage fluid (BALF) samples are standard modalities used to diagnose ventilator-associated pneumonia (45, 46) and can be inundated with pulmonary microbes that grow in cultures, confounding traditional plaque assays (see Fig. S1). As expected from our previous study (9), we found that viral load varied across patient groups where those with severe disease had lower (or absent) viral load at time of sampling (samples were combined with our previously data (9); Fig. 2B). Following 30 min exposure to UVC-irradiation, both virus stock (2.1 × 10^5^ PFU/mL), and respiratory secretions were not detected by plaque assay (Fig. 3C), indicating complete viral inactivation. Additionally, we demonstrate that UVC-inactivation abolishes microbial growth in plaque assays that hampered accurate viral load detection in ETA samples (Fig. S1), which is consistent with established efficacy of UVC-inactivation for a diverse range of pathogenic microbes (47, 48).

**Figure 2.**
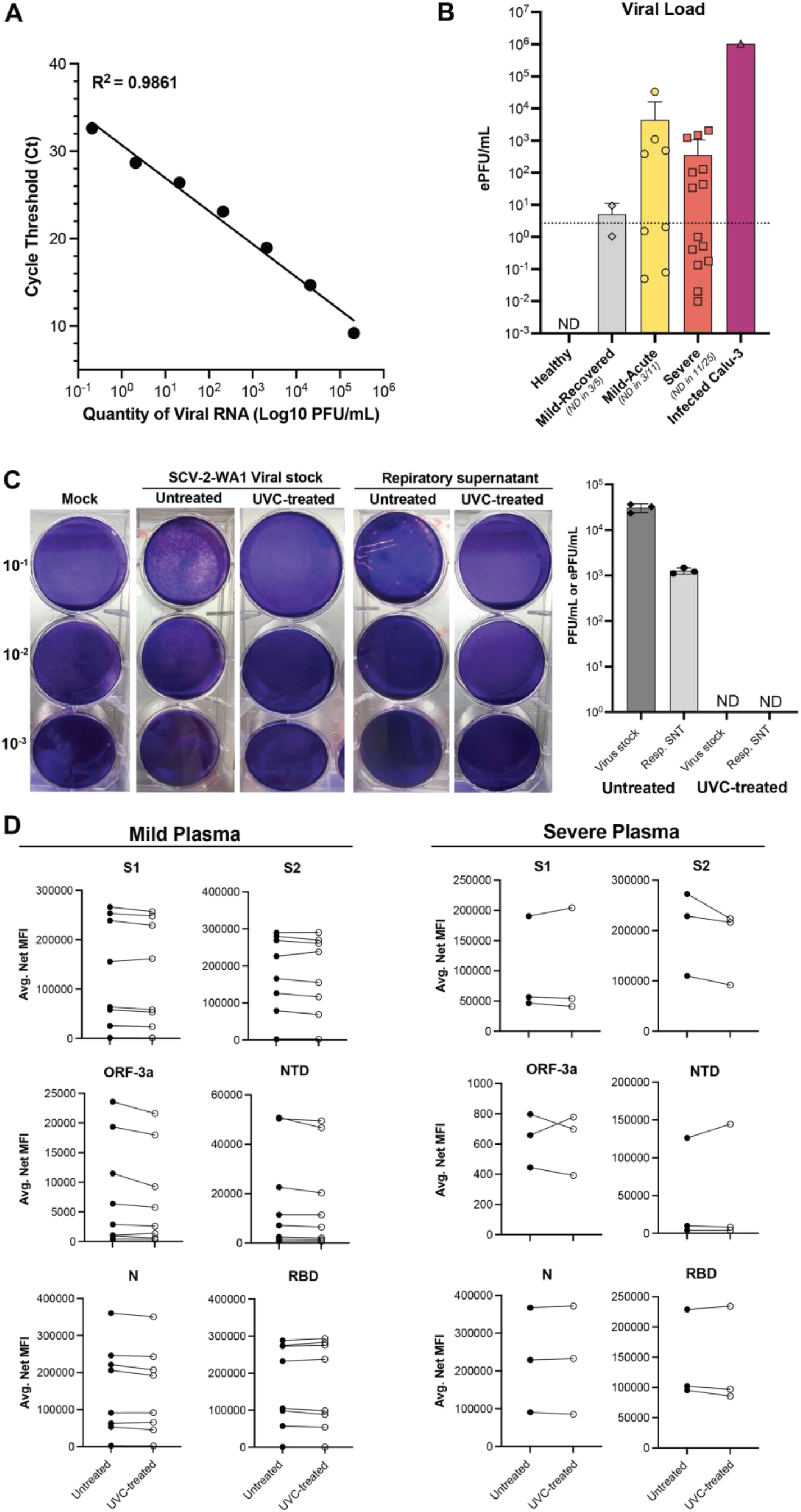
UVC irradiation exposure for 30 minutes inactivates SARS-CoV-2 with minimal effects on antibody/protein detection assays. A. Viral curve generated from serially-diluted SCV2-WA1 stock of a known titer to extrapolate ePFU/mL from RT-qPCR data. B. Viral load (ePFU/mL) in respiratory supernatant (Resp. SNT) from non-induced sputum (healthy and mild) and endotracheal aspirates (ETA; severe) samples using the viral curve generated in A. Dotted line = lower limit of quantification for the ePFU conversion determined by lowest dilution of stock virus (10^−6^) detected by RT-qPCR. ND = Not Detected (by RT-qPCR). C. Representative plaque assays from stock SCV2-WA1 virus and respiratory supernatant (Resp. SNT) samples and quantification of viral load (PFU/mL for SCV2-WA1 stock and ePFU/mL for Resp. SNT from B) before and after UVC-treatment (30 min at ~4000 μwatt/cm^2^). ND = Not Detected (by plaque assay). D. Comparison of SARS-CoV-2 antibody measurements in untreated and UVC-treated plasma samples from mild and severe COVID-19 patients.

**Figure 3.**
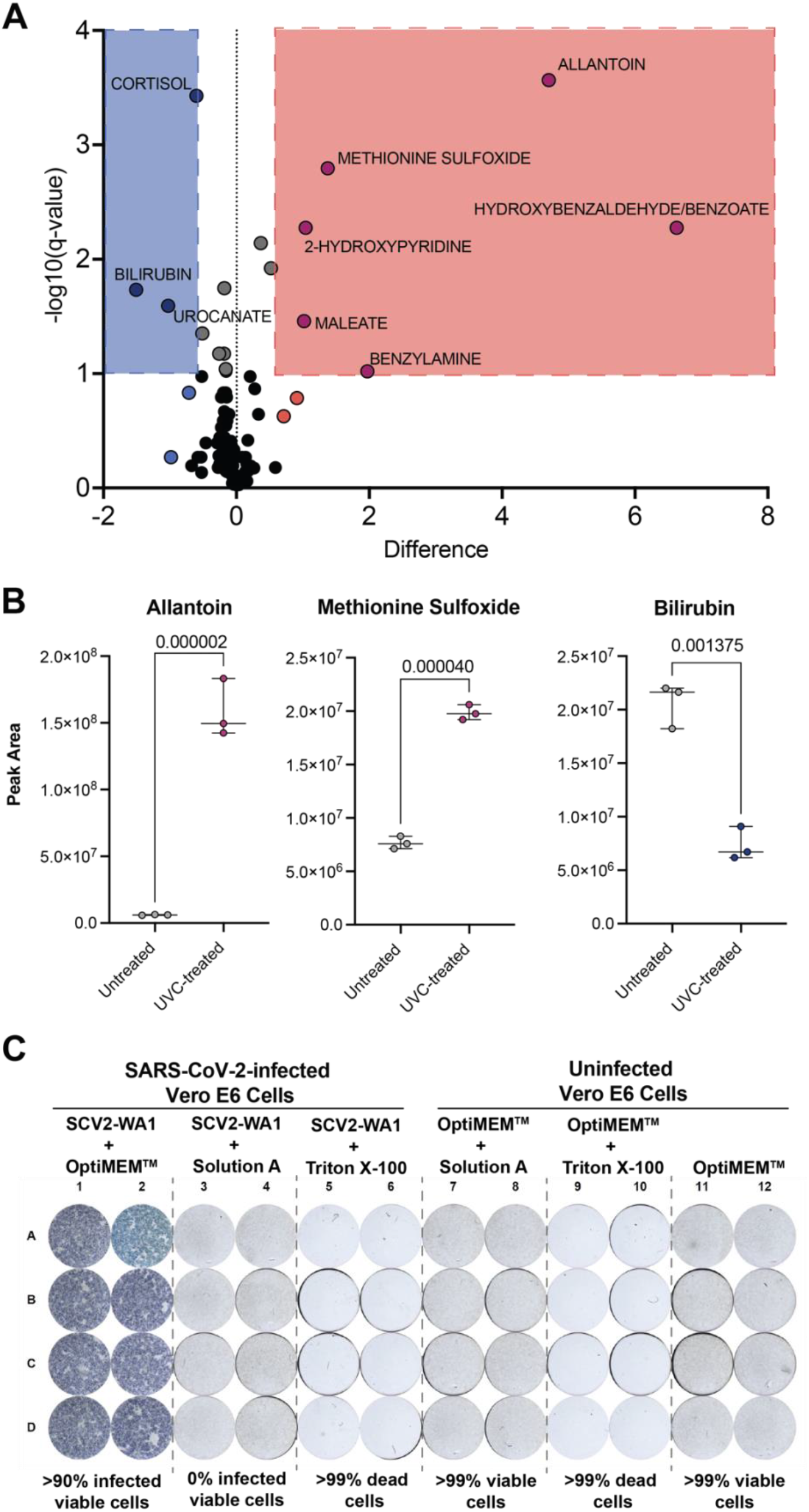
Metabolite extraction solvent (Solution A) completely inactivates SARS-CoV-2 and maintains sample quality for downstream metabolomics assays. A. Volcano plot (FDR <10%) displaying differentially-expressed metabolites in untreated versus UVC-treated NIST (standard) plasma samples. B. Example plots of 3 representatives differentially-expressed, redox-active metabolites in untreated vs. UVC-treated NIST plasma samples. C. FRNA results evaluating inactivation of the metabolite extraction solvent (Solution A) in the standard metabolomic sample processing procedure (see methods) and Triton X-100.

#### 3.2.2 Effects on antibody measurements

To determine the effect of 30 min exposure to UVC-irradiation on protein/antibody detection, we compared untreated and UVC-treated plasma samples from both mild and severe COVID-19 patients (Fig. 2D). We did not observe any significant differences in the levels of anti-SARS-CoV-2 antibodies detected by Luminex proteomic assays. Collectively, these data indicate that 30 min exposure to UVC-irradiation at maximum intensity (~4000 μwatt/cm^2^) is both effective on inactivating high titer stock SARS-CoV-2 and respiratory secretions from COVID-19 patients, with minimal effect on downstream protein/antibody assays.

### 3.3 Inactivation for metabolomics

#### 3.3.1 UVC treatment

To determine optimal inactivation procedure for metabolomics, we first evaluated the effects of UVC-inactivation described above on Standard Reference Material 1950 of metabolites in human plasma (49). UVC-inactivation significantly altered the metabolic profile of samples (Fig. 3A). We show that the differentially-expressed metabolites between untreated and UVC-treated plasma samples are redox active metabolites (Fig. 3B), suggesting that reactive oxygen species (ROS) known to be produced during UVC irradiation (50) lead to sample oxidation during this procedure – similar to ROS oxidation in vivo (51–53). Therefore, UVC-inactivation is not suitable for metabolomic studies, especially if interrogating redox active metabolites.

#### 3.3.2 Treatment with organic solvents

We then wanted to test if the standard metabolomic sample extraction procedure with organic solvents (see methods), could successfully inactivate SARS-CoV-2-infected non-cellular products (such as respiratory secretions or plasma) as an alternative to UVC-inactivation. Specifically, we used a 1:1 mixture of acetonitrile and methanol including a deuterated internal standard, administered at 4 volumes relative to starting sample. We performed FRNAs with stock SCV2-WA1 (untreated), and virus treated with either the extraction solvent or Triton X-100, which has been shown to inactivate SARS-CoV-2 (17, 18). Incubation of SCV2-WA1 virus stock 1:4 with the extraction solvent was sufficient to fully inactivate virus along with Triton X-100 (Fig. 3C). Cells incubated with the extraction solvent remained viable (Fig. 3C), confirming virus inactivation independent of Vero E6 cytotoxicity, which was observed for other chemical reagents/surfactant such as Triton X-100 (Fig. 3C). Therefore, these results demonstrate that the standard metabolomic assay sample processing procedure with 4 volumes of our extraction solvent is sufficient to inactivate SARS-CoV-2-infected non-cellular samples while retaining sample integrity as compared to UVC-inactivation.

### 3.4 Inactivation for scRNA-seq

To better understand efficacy of SARS-CoV-2 inactivation in droplet-based scRNA-seq pipelines—specifically the 10X Genomics platform—we evaluated viral inactivation at the two early steps in the manufacturer’s instructions. First, we evaluated the reagent cytotoxicity of the 10X Genomics encapsulation emulsion on Vero E6 cells by performing plaque assays with the emulsion free of encapsulated cells. We demonstrate that the reagents in the emulsion (many of which are proprietary) are not inherently cytotoxic to Vero E6 monolayers, allowing us to evaluate viral inactivation by standard plaque assay (Fig.4a).

**Figure 4.**
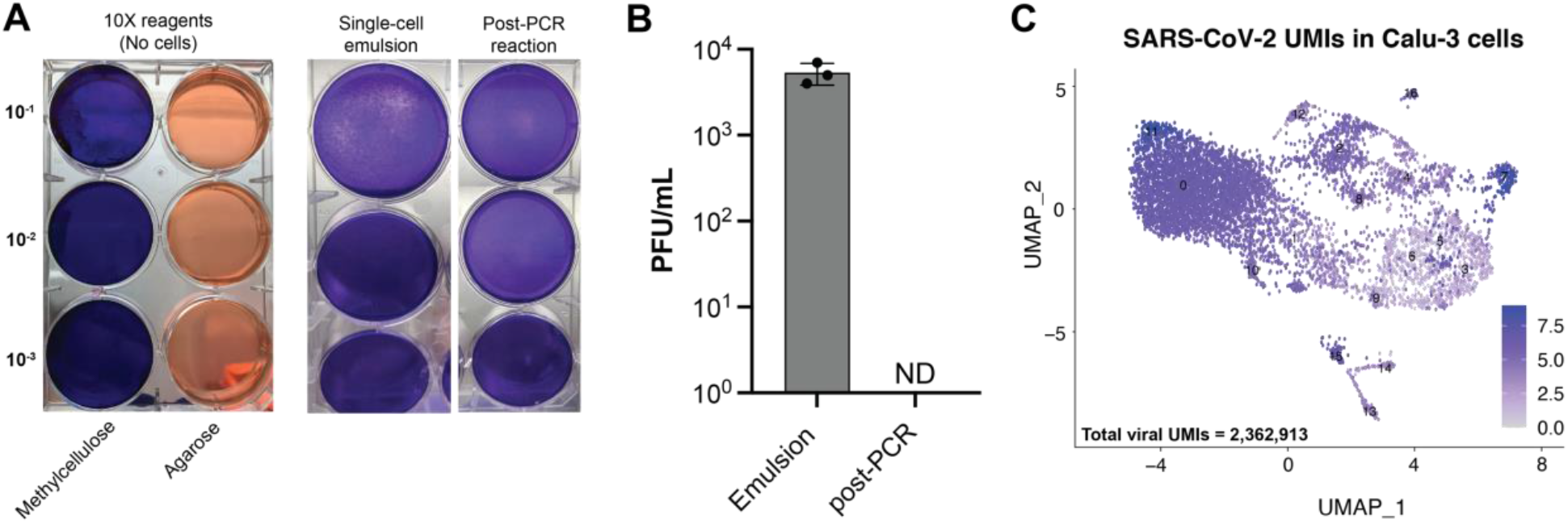
Heat inactivation during cDNA synthesis completely inactivates SARS-CoV-2 in scRNA-seq emulsions. A. Representative plaque assays performed using 10X Genomics’ emulsion reagents alone (without cells) to evaluate reagent cytotoxicity on Vero E6 cells and single-cell emulsion with SCV2-WA1-infected Vero E6 cells (MOI 0.04), and the same emulsion after cDNA synthesis PCR reaction (45 min at 53°C followed by 5 min at 85°C). B. Quantification of viral load in single-cell emulsions of SCV2-WA1-infected Calu-3 cells (MOI 0.04) immediately after encapsulation and following PCR reaction for cDNA synthesis. N=3 independent samples. ND = Not Detected (by plaque assay). C. UMAP visualization of scRNA-seq data from SCV2-WA1-infected Calu-3 cells (MOI 0.04; N=8061 cells) showing expression of the 12 SARS-CoV-2 genes and total viral UMIs (inset).

We next evaluated the efficacy of viral inactivation following single-cell encapsulation (which contains a proprietary lysis solution) of SCV2-WA1-infected (MOI 0.04) Vero E6 or Calu-3 cells. We show that the standard single-cell encapsulation step alone is not sufficient to fully inactivate SARS-CoV-2, which could be detected in subsequent plaque assays (Fig. 4A,B). Therefore, we tested the efficacy of the first round cDNA synthesis reaction, which includes exposure to temperatures ≥53°C, in viral inactivation. We show that after the PCR reaction (45 min at 53°C followed by 5 min at 85°C), infectious SARS-CoV-2 was not detectable by plaque assay (Fig. 4A,B). Taken together, our data demonstrate that scRNA-seq emulsions of SARS-CoV-2-infected cells are fully inactivated only after heat inactivation following the cDNA synthesis reaction using the conditions described in the manufacturer’s protocol (45 min at 53°C followed by 5 min at 85°C), which can then be transferred to lower containment for library preparation and sequencing. Indeed, after sequencing, we find >2.3 million viral transcripts (or unique molecular identifiers; UMI) in ~8,000 SCV2-WA1-infected Calu-3 cells (Fig. 4C).

## 4. Discussion

To date, there have been multiple studies to evaluate efficacy of viral inactivation procedures using heat, chemicals, and UVC irradiation on SARS-CoV-2-infected samples (14–18). Many of these studies have evaluated traditional procedures established for SARS-CoV and MERS-CoV and were found to have comparable efficacies (12, 13, 20, 21). Here, we add to these studies by evaluating inactivation procedures performed under optimized conditions that allow for downstream processing/analysis for contemporary immunology assays with limited effects on assay readouts. We caution that all procedures are performed under the specified conditions and those that differ from what have been described here should be evaluated on viral stocks and patient samples before transferring to lower biosafety containment.

Since its induction in the 1960s, flow cytometry has been the preeminent technology for single-cell analysis (54) particularly for investigating the heterogeneity of the immune system in health and disease (55, 56). Indeed, Hi-D flow cytometry has been a pivotal tool in dissecting the complex immunophenotypes of leukocytes in COVID-19 (9, 29, 30, 57, 58). We evaluated the ability of commercially-available fixatives commonly used in flow cytometry (formaldehyde-based) to fully inactivate SARS-CoV-2-infected cells to facilitate transfer of cells from BSL3 to BSL2 for data acquisition (9). Here, we show that treatment with 4% PFA or 1.6X BD FACS™ Lysis solution for 30 min at RT was sufficient to completely inactivate SARS-CoV-2-infected cells, even at viral titers higher than in cells from infected patients. Therefore, most common fixation protocols and reagents (e.g., BD Cytofix/Cytoperm™, BioLegend^®^ Fixation Buffer, etc.) that contain ≥4% PFA are suitable for preparing fluorescently-stained, SARS-CoV-2-infected cells for transfer out of BSL3 containment after 30 min exposure. Conversely, lower concentrations of PFA (i.e., 2%) required longer exposure time (at least 60 min at RT) to fully inactivate samples.

Similarly, we show that UVC irradiation (~4000 μwatts/cm^2^) for 30 min is sufficient to fully inactivate high titer SCV2-WA1 viral stocks and respiratory supernatants from patients with minimal effects on protein/antibody detection (59), which will promote further studies on secretions from ETA and/or BALF to better understand local versus systemic responses (9, 25). It is important to note that a previous study on UVC-inactivation of SARS-CoV found BSA to protect virus from UVC-inactivation even after 60 min exposure (20). Therefore, we avoided BSA in solutions used in respiratory sample preparation (9), and plasma samples were inactivated prior to dilution in the Luminex proteomic assay buffer (1% BSA in 1X PBS, see methods).

Despite having negligible effects in the proteomic assays, we did observe that UVC inactivation significantly altered metabolomic profiles is human plasma samples. Specifically, we show that redox active metabolites such as methionine and urate are oxidized following UVC inactivation, significantly increasing signals for methionine sulfoxide and allantoin, respectively (51, 52). Similarly, bilirubin, which is oxidized to biliverdin, is significantly decreased with UVC treatment (53). Therefore, UVC-inactivation of clinical samples could lead to misleading biological interpretations, artificially skewing sample metabolites to a more oxidized profile (51–53). However, high concentration methanol (≥80%) (60, 61) and methanol/acetone mixtures (13, 22) have previously been shown to successfully inactivate many viral infected samples including SARS-CoV-2 (17–19). The extraction solvent we used is similar to that used in many metabolomic sample preparation techniques, and was sufficient to inactivate virus while maintaining data integrity.

Systems immunology approaches, including multi-omic scRNA-seq, have greatly advanced our understanding of COVID-19 immunity and pathogenesis (9, 31, 32). However, a detailed report on inactivation efficacy of scRNA-seq pipelines is notably lacking. We demonstrate that in the standard 10X Genomics pipeline, encapsulation alone was insufficient to fully inactivate virus. According to the manufacturer’s guidelines, a cDNA synthesis reaction is the next immediate step after encapsulation (see methods). Though many studies have evaluated the efficacy of heat inactivation for SARS-CoV-2 and demonstrated 45 min at 56°C and 5 min at 100°C are sufficient to fully inactivate virus (14–19), none have tested the specific conditions for the cDNA synthesis reaction (45 min at 53°C followed by 5 min at 85°C). Here, we expand on the previous studies by demonstrating that the cDNA synthesis reaction in the standard 10X Genomics pipeline successfully inactivates SARS-CoV-2 and allows for transfer to lower containment for subsequent processing and library generation procedures.

Thus, we report optimized methods of viral inactivation that have minimal, if any, adverse impact on immunological studies of infected culture-derived and patient samples, permitting safe transfer to lower containment laboratories (BSL2) for final processing and data acquisition. Taken together, this suite of inactivation procedures can serve as guidelines for rapid initiation of research as the COVID-19 pandemic continues.

## Acknowledgements

This work was supported by funds from NIH/NIAID R01AI123126 (EEBG) and R01AI123126-05S1 (EEBG), NIH T32-HL116271-07 (RPR), the Program for Breakthrough Biomedical Research, and Lowance Center for Human Immunology (EEBG), the NSF EAGER award 2032273 (RT, RFS, KZ), and the Woodruff Health Science Center COVID-19 CURE award (RT, RFS, KZ). DJE was supported, in part, by the Laney Graduate School Fellowship (Emory) and JDC was supported, in part, by CF@LANTA, a component of Emory University and Children’s Healthcare of Atlanta. The funders had no role in study design, data collection and analysis, decision to publish, or preparation of the manuscript. We thank Fred Souret (10X Genomics) for providing additional reaction kits to test viral inactivation in scRNA-seq assays, Doan Nguyen for technical direction with the serology experiments, and the Emory Pediatric/Winship Flow Cytometry Core (access supported in part by Children’s Healthcare of Atlanta) for their support with flow cytometry experiments. We thank Rafi Ahmed (Emory) and Jacob Kohlmeier (Emory) for kindly providing the Vero E6 and Calu-3 cell lines, respectively. We thank Ann Chahroudi, Nils Schoof, Kira Moresco, and Stacy Heilman of the Department of Pediatrics (Emory), along with the Emory Biosafety Officers Kalpana Rengarajan and Esmeralda Meyer and the Institutional Biosafety Committee (IBC) for their assistance with the BSL3 facility and protocol review/approval.

## Declaration of interest

FEL is the founder of MicroB-plex, Inc., serves on the SAB of Be Bio Pharma, receives grants from BMGF and Genentech, and receives royalties from BLI, inc. All other authors have no competing interest to declare.

## Author contributions

**Devon J. Eddins**: Conceptualization, Data curation, Formal analysis, Investigation, Methodology, Validation, Visualization, Writing - original draft, Revising & editing – final draft. **Leda Bassit**: Investigation, Methodology, Data curation, Formal Analysis, Visualization. **Joshua D. Chandler**: Investigation, Methodology,Data curation, Formal Analysis, Visualization. **Natalie S. Haddad** Investigation, Data curation, Formal Analysis. **Katie L. Musall**: Investigation. **Junkai Yang**: Formal Analysis, Visualization. **Astrid Kosters**: Investigation. **Brian S. Dobosh**: Investigation, Formal Analysis. **Mindy R. Hernández**: Resources. **Richard P. Ramonell**: Resources. **Rabindra M. Tirouvanziam**: Supervision, Resources, Data curation. **F. Eun-Hyung Lee**: Supervision, Resources, Data curation. **Keivan Zandi**: Resources, Investigation, Methodology, Data curation, Formal analysis, Supervision. **Raymond F. Schinazi**: Supervision, Resources, Data curation. **Eliver E.B. Ghosn**: Conceptualization, Data curation, Formal analysis, Funding acquisition, Methodology, Project administration, Resources, Supervision, Visualization, Writing – original draft; Revising & editing – final draft. All authors discussed the results and read and approved the final manuscript.

---

**Figure S1.**
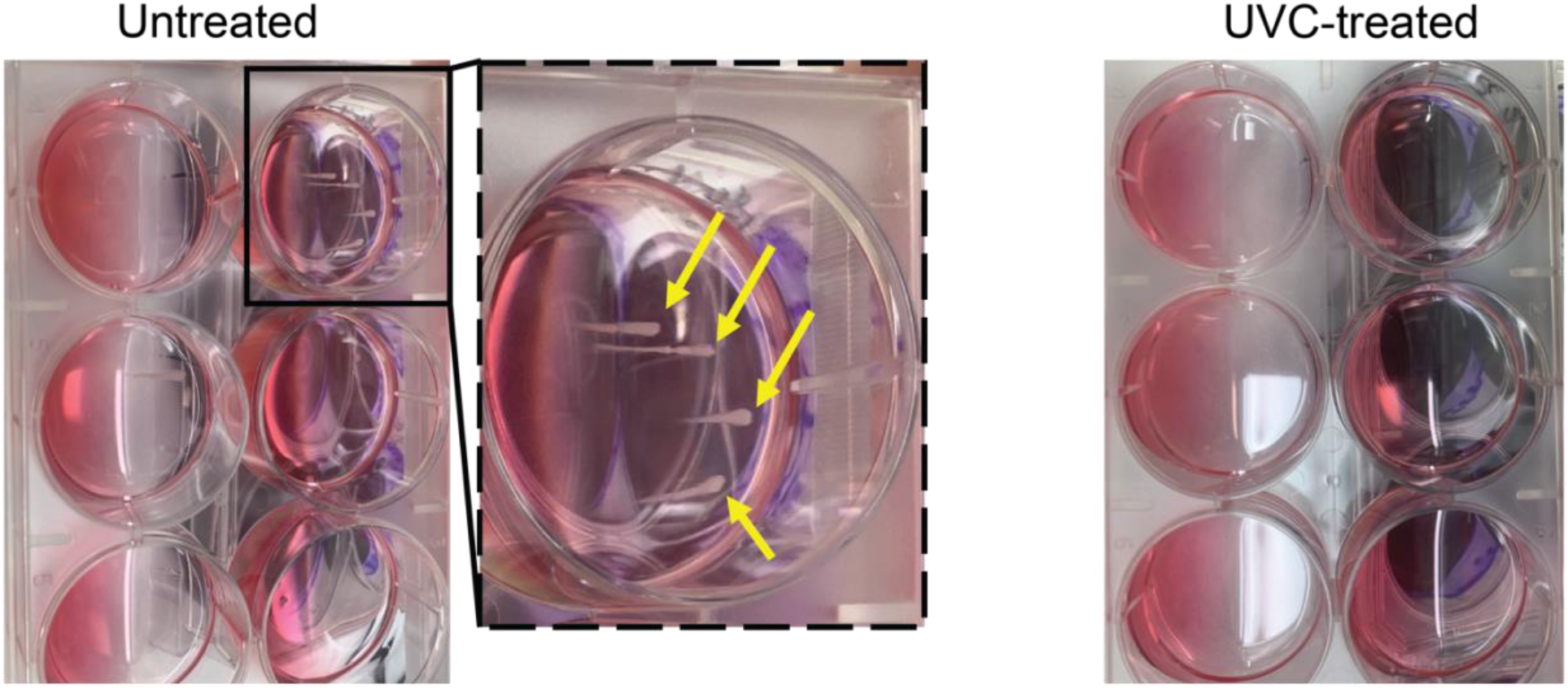
UVC inactivation abolishes microbial growth in plaque assay cultures. Representative image of plaque assay cultures for respiratory supernatant samples (severe COVID-19 patients) before and after UVC-treatment (30 min at ~4000 μwatt/cm^2^). Yellow arrows indicate microbial growth.

## Notes

### Competing Interest Statement

The authors have declared no competing interest.

